# SequelTools: A Suite of Tools for Working with PacBio Sequel Raw Sequence Data

**DOI:** 10.1101/611814

**Authors:** David E. Hufnagel, Matthew B. Hufford, Arun S. Seetharam

**Affiliations:** Department of Ecology, Evolution and Organismal Biology, Iowa State University, Ames, IA, 50011, USA; Genome Informatics Facility, Iowa State University, Ames, IA, 50011, USA

## Abstract

**Background:** PacBio sequencing is an incredibly valuable third-generation DNA sequencing method due to very long read lengths, ability to detect methylated bases, and its real-time sequencing methodology. Yet, hitherto no tool was available for analyzing the quality of, subsampling, and filtering PacBio data.

**Results:** Here we present *SequelTools*, a command-line program containing three tools: Quality Control, Read Subsampling, and Read Filtering. The Quality Control tool quickly processes PacBio Sequel raw sequence data from multiple SMRTcells producing multiple statistics and publication-quality plots describing the quality of the data including N50, read length and count statistics, PSR, and ZOR. The Read Subsampling tool allows the user to subsample reads by one or more of the following criteria: longest subreads per CLR or random CLR selection. The Read Filtering tool provides options for normalizing data by filtering out certain low-quality scraps reads and/or by minimum CLR length. *SequelTools* is implemented in bash, R, and Python using only standard libraries and packages and is platform independent.

**Conclusions:** *SequelTools* is a program that provides the only free, fast, and easy-to-use quality control tool, and the only program providing this kind of read sumbsampling and read filtering for PacBio Sequel raw sequence data, and is available at https://github.com/ISUgenomics/SequelTools

## 1 Background

The third-generation of sequencing is here and making tremendous impact in the field of genomics. The primary contenders in third-generation sequencing are Pacific Biosciences (PacBio) (Sequel, Sequel2) and Oxford Nanopore (MinION, GridION, and PromethION). These new sequencing platforms are undergoing active development and pushing boundaries in terms of total output, read length, sequencing time, cost reduction and read accuracy (Schadt *et al.*, 2010; Goodwin *et al.*, 2016). Recently introduced PacBio Sequel/Sequel2 platforms, which rely on Single-Molecule Real Time (SMRT) sequencing technology, are one of the most widely used long-read sequencing approaches (Rhoads and Au, 2015; Goodwin *et al.*, 2016). In contrast to second-generation methodologies, PacBio provides longer length reads, in much less time, with greatly reduced-content bias, and an ability to distinguish between methylated and unmethylated bases (Schadt *et al.*, 2010; Ross *et al.*, 2013; Rhoads and Au, 2015; Goodwin *et al.*, 2016). Accuracy has also substantially improved from previous long-read platforms.

Similar to the previous RSII platform, Pacbio Sequel uses the SMRTBell, a double stranded DNA molecule that loops around the ends, as the template for sequencing. The polymerase runs through the template continuously, sequencing the DNA by adding nucleotides in both the forward and reverse orientation. The contiguous sequence generated by the polymerase during sequencing is referred to as a “polymerase read” or a Continuous Long Read (CLR). This CLR read may include sequence from adapters and multiple copies of inserts, because it traverses the circular template many times. The CLRs are processed to remove adapter sequences and to retain only the insert sequence, called “subreads”. Multiple copies of subreads generated from the single SMRTBell can then be collapsed to a single, high-quality sequence, called the “read of insert” or Circular Consensus Sequence (CCS) (Rhoads and Au, 2015; Ardui *et al.*, 2018).

Sequencing is performed within a SMRTcell which contains tens of thousands of zero-mode waveguides (ZMWs). These ZMWs contain a light-detection module and an immobilzed polymerase enzyme. The template (SMRTBell) is introduced into the ZMW and nucleotide bases labeled with different fluorophores are sequentially added. Each base incorporation will result in the release of a fluorophore, producing distinct light wavelengths per base. All ZMWs within a SMRTcell are processed in parallel, sequencing thousands of templates at the same time. Thus the number of productive ZMWs (ZMWs that received exactly one template) will directly indicate the productivity of the SMRTcell. The length distribution for the polymerase reads/subreads also provides a useful metric of run quality (Ardui *et al.*, 2018).

When working with sequence data it is important to be aware of sequence quality before using the data for downstream analysis, otherwise poor quality reads could lead to spurious results. Programs or web applications that accomplish this task are often referred to as quality control (QC) tools. One popular QC tool for short-read sequence is *fastQC* (Andrews *et al.*, 2012). *fastQC* works well for short reads, but is not appropriate for long reads found in third-generation sequences. Hitherto, there are no freely available tools for assessing the quality of raw PacBio sequence data. The development of a fast, free, and easy-to-install and use program to assess raw sequence quality is therefore crucial for any scientist making use of PacBio Sequel sequence data. There is also no known tool that allows users to subsample reads to reduce data size for testing or filter reads so as to normalize the data.

Here we present *SequelTools*, an efficient and user-friendly program with multiple tools including a QC tool that calculates multiple standardized statistics and creates publication-quality plots describing the quality of raw PacBio Sequel data, a Read Subsampling tool that allows the user to subsample their data by either longest subreads per CLR and/or random CLRs, and a Read Filtering tool that filters the user’s data by one or more chosen criteria. In conclusion, *SequelTools* provides essential functions for analyzing and processing Pacbio Sequel raw sequence data, and is the only program to date of its kind. We believe *SequelTools* will therefore facilitate the use of PacBio sequence data for researchers in a variety of disciplines.

## 2 Implementation

*SequelTools* uses only standard libraries and packages within bash, R and Python in order to facilitate quality assessment, data filtering and normalization of raw PacBio Sequel data. While it can be run on a single SMRTcell, *SequelTools* is designed to run across multiple SMRTcells simultaneously. The main script is written in bash which calls Samtools for converting between BAM (Binary Alignment/Map format) and SAM (Sequence Alignment/Map format) format, Python for calculations, and R for plotting. Python 2 or 3 can be used, and the version is determined automatically by the program. *SequelTools* is fast (with the exception of subsampling for longest subreads), easy to use, and works on any operating system from the command-line. *SequelTools*, in its current form, is composed of three “tools”, which can be used one at a time using regular command-line arguments. The three tools implemented in this program are 1) Quality Control, 2) Read Subsampling, and 3) Read Filtering. *SequelTools* uses BAM format files as input because raw Pacbio Sequel sequence files come in BAM format. PacBio sequence files include both subreads files containing reads of interest and scraps files with additional reads generated during the sequencing process. For all of *SequelTools*’ tools subreads files are required, and for some scraps files are also required.

### 2.1 The Quality Control Algorithm

One of *SequelTools*’ tools is the QC tool. This tool creates tables and plots summarizing the quality of PacBio Sequel data. The QC tool does not require scraps files. With scraps files, the QC tool takes longer to run, but also produces additional plots and provides more information in standard plots using additional information concerning CLRs. The QC tool calls Samtools (Li *et al.*, 2009) and awk to convert BAM files to SAM format and extracts only needed information (Figure 1). Then Python is used to make all necessary calculations, producing intermediate data files that are passed to R. Note that when including scraps files only normal scraps reads are used for downstream QC analyses. By default these intermediate data files will be deleted at the end of the program’s operation, but they will be retained if the user selects the appropriate arguments. At this point, reads are organized into up to four read groups: 1) subreads, 2) longest subreads, 3) CLRs, and 4) subedCLRs (CLRs containing subreads). When scraps are included, the default is to use all four read groups, but the user can request only two groups if preferred: 1) subreads and 2) subedCLRs. Alternatively, if scraps are excluded the two read groups are 1) subreads and 2) longest subreads.

**Figure 1:**
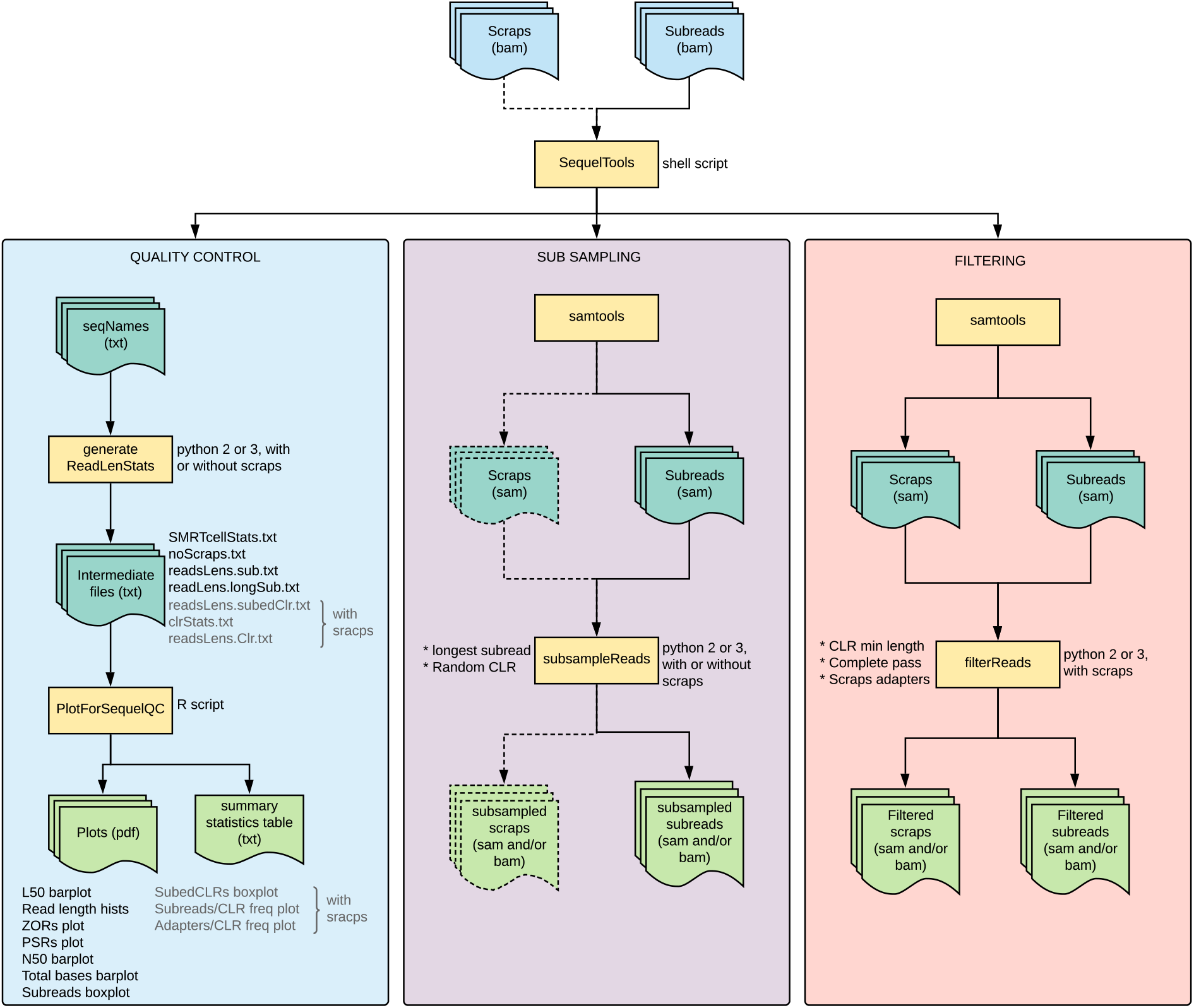
*SequelTools* flowchart. A flowchart of how SequelTools processes input files and uses them to perform QC, Read Subsampling and Read Filtering functions.

Final plots and tables are produced in R, including a tab-delimited table of summary statistics, which can be viewed easily in Microsoft Excel, as well as several publication-quality PDF (Portable Document Format) plots. A subset of these plots can be seen in Figure 2; All plots can be seen in Figure 4 The summary statistics table includes information for all chosen read groups for each SMRTcell. Statistics include number of reads, total bases, mean and median read length, N50, L50, PSR, and ZOR. PSR is the polymerase-to-subread ratio and is calculated as follows: total bases from the longest subreads per CLR divided by the total bases from subreads. ZOR is the ZMW occupancy ratio and is calculated as the number of CLRs with subreads divided by the number of subreads.

**Figure 2:**
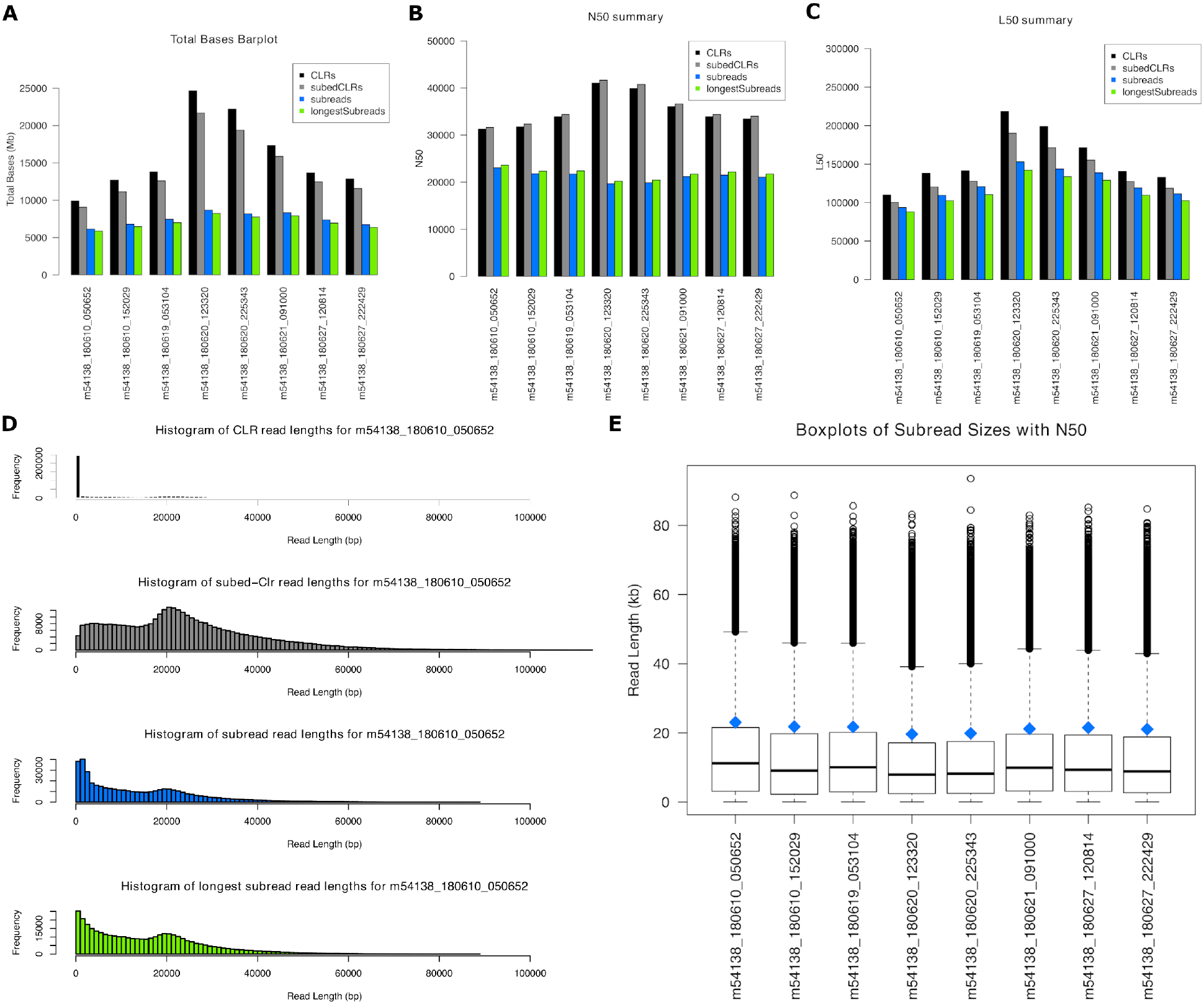
Selected QC plots generated by *SequelTools*: (A) A barplot of the sum of read lengths, (B) A barplot of N50s, (C) A barplot of L50s, (D) Read length histograms for m54138 180610 050652, (E) and Boxplots of subread lengths with N50s as blue diamonds, for all SMRTcells in our benchmarking data set.

### 2.2 The Read Subsampling Algorithm

Another of *SequelTools*’ tools is the Read Subsampling tool. This tool allows the user to subsample their BAM format sequence files by longest subreads per CLR or random CLR selection. Subsampling by the longest subreads per CLR simply creates a new sequence file for each SMRTcell containing only the longest subread for each CLR. Similarly, subsampling by random CLR creates a new sequence file for each SMRTcell containing only reads associated with randomly selected CLRs. The Read Subsampling tool does not require scraps files, but can take advantage of them, if provided by the user, when subsampling for randomly selected CLRs. When subsampling by longest subreads alone, if scraps files are provided which is not recommended, a new subsampled scraps file will be generated identical to the original scraps file. As with the QC tool, using scraps files will increase *SequelTools*’ runtime, mostly due to the need to convert additional BAM files to SAM format.

The Read Subsampling tool first converts BAM sequence files to SAM format using Samtools (Figure 1). Next the SAM files are processed using Python. Python parses through the SAM files and saves necessary information. When subsampling by longest subreads coordinate information associated with CLR IDs are saved by Python, and when subsampling by random CLRs all CLR IDs are saved by Python. Subsampling CLR IDs and/or read IDs based on chosen criteria comes next, followed by parsing input SAM files again and outputting all reads associated with subsampled longest subreads and/or randomly subsampled CLRs in the SAM format. Finally, if desired, the Read Subsampling tool will convert the output files to BAM format using Samtools.

### 2.3 The Read Filtering Algorithm

An additional tool available via *SequelTools* is the Read Filtering tool. This tool provides the user a filtering functionality and requires both scraps files and subreads files to function. Filtering can be done using one or more of the following criteria: 1) minimum CLR length, 2) having at least one complete pass of the DNA molecule through the polymerase, or 3) normal adapters for scraps. The latter two criteria apply only to scraps files, and are recommended for most downstream analyses involving scraps files. Filtering by minimum CLR length takes input BAM format subreads and scraps files and yields files with only CLRs having a total length, including all provided scraps, greater or equal to the threshold value provided by the user. Filtering by complete passes of the DNA molecule takes input BAM format subreads and scraps files and creates an output containing only scraps reads with at least one full pass of the DNA molecule through the polymerase. Filtering by normal scraps adapters takes BAM format subreads and scraps files as input and creates an output with only normal scraps adapters, defined as having a ZMW classification annotation of ‘N’ for ‘normal’ and a scrap region-type annotation of ‘A’ for ‘adapter’ (Biosciences, 2019a).

The Read Filtering tool starts by using Samtools to convert BAM sequence files to SAM format (Figure 1). Then the SAM files are processed with a Python script. Python first extracts coordinates from scraps and subreads files. Next, if filtering by CLR length, CLRs are assembled from subreads and scraps coordinate data, CLR lengths are calculated, and CLR IDs are stored that do not pass the minimum length threshold. If filtering by CLR length the subreads input file is iterated through again and only reads containing CLRs that pass the minimum length threshold are written to a new output file. If filtering by either the number of passes of the DNA molecule or normal scraps adapters, the scraps file is then iterated through again and only reads containing information indicating they pass all chosen scraps thresholds are written to a new output file. Lastly, upon the user’s request, the Read Filtering tool will convert the output files to BAM format using Samtools.

### 2.4 Using The Quality Control Tool

The QC tool requires only two arguments: ‘-u’ and ‘-t’. In its most basic form with scraps files it looks like so:

~~~
bash SequelTools.sh -t Q -u subFiles.txt -c scrFiles.txt

or without scraps files:

bash SequelTools.sh -t Q -u subFiles.txt
~~~

Where the ‘-t’ argument tells *SequelTools* which tool to use, in this case ‘Q’ for the Quality Control tool and subFiles.txt and scrFiles.txt are file-of-filename files containing the names of subreads and scraps BAM files, respectively, with one name per line. While ‘-u’ and ‘-t’ are the only required arguments, the user can choose from other arguments including the number of threads to use for running Samtools, whether to keep intermediate files, and how many read groups and plots are desired. The summary statistics table as well as all plots, except the read length histograms and frequency plots, present the data from all SMRTcells together. For read length histograms and frequency plots, separate files are generated for each SMRTcell with either one plot per group for histograms or one plot total for frequency plots.

While the summary statistics table is always produced, the user can request more or fewer plots based on their needs. The full suite of plots with scraps files includes barplots of A) N50s, B) L50s, C) total bases, and D) read length; frequency plots of E) subreads per subedCLR, and F) adapters per CLR; boxplots of G) subread lengths and H) subedCLR lengths; and I) ZOR and J) PSR plots (Figure 4). The user can also request an intermediate (A,C,G,H,I, & J) or basic (A & C) suite of plots. Without scraps files, the full suite of plots is A,B,C,D,G,I, & J, the intermediate collection is A,C,G,I, & J, and the basic set is A & C (Figure 1). With or without scraps the intermediate selection of plots is default.

Some users may want to modify the provided R script to change plot format or to make entirely new plots. In the case that a user wishes to use a custom R script for plotting, we recommend the user run the QC tool once using the ‘-k’ argument to generate the intermediate data files and retain them at the end of the QC tool’s operation. Next, the user will need to create their custom R script. If the user wishes to modify *SequelTools*’s QC plots we recommend the user start by copying and renaming either ‘plotForSequelTools wScraps.R’ or ‘plotForSequelTools noScraps.R’ depending on whether the user is running the QC tool with or without scraps files, respectively. To aid in the process of testing the custom R plotting script we have added an argument ‘-s’ which will skip the read length calculations with Samtools and the statistical calculations with Python which, together, generate the intermediate data files. Together these steps make up most of the runtime of the QC tool, therefore skipping these steps allows for rapid testing of an alternative plotting script. Whether the user is modifying a *SequelTools* R plotting script or using one created from scratch the user will need to provide the custom plotting script to *SequelTools*. This can be done using the ‘-r’ argument followed by the name of the custom script. When testing the custom R script the ‘-k’ argument will remain necessary, otherwise the intermediate files will all be deleted at the end of the QC tool’s operation. Altogether, running the QC tool with an alternative R plotting script with minimal recommended arguments looks like so with scraps files:

~~~
bash SequelTools.sh -u subFiles.txt -c scrFiles -k -s -r altRscript wScraps.R

or without scraps files:

bash SequelTools.sh -u subFiles.txt -k -s -r altRscript noScraps.R
~~~

### 2.5 Using The Read Subsampling Tool

The Read Subsampling tool requires four arguments: ‘-t’, ‘-u’, ‘-c’, and ‘-T’. In its simplest construction, subsampling using both criteria, it looks like this with scraps files:

~~~
bash SequelTools.sh -t S -u subFiles.txt -c scrapsFiles.txt -T lr

or without scraps files:

bash SequelTools.sh -t S -u subFiles.txt -T lr
~~~

The ‘-t’,‘-u’, and ‘-c’ arguments have been explained previously in this implementation section. The ‘-T’ argument allows the user to control which of the two possible criteria with which to subsample. Just like with the QC tool, with the Read Subsampling tool the user may control the number of threads used when running Samtools, using the ‘-n’ argument. When subsampling randomly by CLR the user will likely wish to provide the argument ‘-R’, which specifies what proportion of CLRs to be retained in that process. We have also designed an argument, ‘-f’, which allows the user to decide whether they want the output of the Read Subsampling tool to be in BAM, SAM or both formats.

### 2.6 Using The Read Filtering Tool

The Read Filtering tool requires four arguments, ‘-t’, ‘-u’, ‘-c’ and at least one of the three filtering criteria arguments ‘-C’, ‘-P’, or ‘-N’. This means that scraps files are required in addition to subreads files. The ‘-t’, ‘-u’, and ‘-c’ arguments have been explained previously in this implementation section. The ‘-C’ argument tells the Read Filtering tool to filter by minimum CLR length. The ‘-P’ argument tells the Read Filtering tool to filter by the number of complete passes of the DNA template. The ‘-N’ argument tells the Read Filtering tool to filter by normal scraps adapters. When using the ‘-C’ argument, the ‘-Z’ argument is also required, which sets the minimum CLR length for filtering. Keep in mind when running the CLR length filter the calculated CLR length will include all subreads and scraps data provided including scraps reads with fewer than one complete pass of the DNA molecule by the polymerase and scraps that do not pass the normal adapter scraps filter. This remains true when all three filters are run simultaneously. If the user wishes for CLR length to be calculated for the purpose of applying a minimum length threshold to include scraps reads, but to exclude these less desirable scraps reads, they must run the filters for number of complete passes of the DNA molecule and normal adapter scraps reads first and then run the filter for minimum CLR length using BAM files generated by the Read Filtering tool in the first run.

Here is an example of running the Read Filtering tool of *SequelTools*, in its simplest construction, using all three possible filtering criteria and using 1000 base pairs as the minimum CLR length:

~~~
bash SequelTools.sh -t F -u smallSubs.txt -c smallScraps.txt -C -P -N -Z 1000
~~~

As with *SequelTools*’ other tools, the ‘-n’ argument can be used to set the number of threads Samtools uses when converting between BAM and SAM format. Also, as with the Read Subsampling tool the ‘-f’ argument can be used to set the output format as BAM, SAM, or both.

## 3 Results/Discussion

### 3.1 The Quality Control Tool

It is of great importance that a determination of the quality of sequence data be made before said sequence is used for any downstream analysis. Programs like fastQC (Andrews *et al.*, 2012) are widely used for short reads, but do not provide all the necessary metrics necessary for quality control of long read sequences like those created by the PacBio Sequel system. While a quality control tool for Oxford Nanopore Technologies’ MinION seqence is available (Lanfear *et al.*, 2019), there is no such tool available for the newest sequencing technology from PacBio, PacBio Sequel. Due to improvements in data formats and the technology itself, previous base quality programs for PacBio RSII (Desvillechabrol *et al.*, 2018; mhsieh, 2019) data are no longer valid for assessing the quality of PacBio Sequel data. Currently, the only program that provides quality assessment for PacBio Sequel raw sequence data is the instrumentation software itself, SMRT Link, a linux-only, computationally intensive webtool where the user must upload their data files one at a time. Furthermore, SMRT Link can only be installed by root users, requiring the installation of 23 external programs to run, and generates non-downloadable plots after setting up a web server (Biosciences, 2019c). In fact, the difficulty of using SMRT Link for high-throughput data was the motivation for writing *SequelTools*. Even for users who have the specialized skills required to install and run SMRT Link, running the QC tool via *SequelTools* would be much faster and simpler, free, platform independent, and would produce publication-quality plots.

### 3.2 The Read Subsampling Tool

In addition to the QC tool, a Read Filtering tool has been implemented in *SequelTools*, which can subsample longest subreads per CLR or randomly selected CLRs. When PacBio sequence is generated, the DNA template is often sequenced many times resulting in several copies of the sequence of interest and multiple subreads per CLR. One effective approach for handling these multiple subreads is to generate a consensus sequence (Wenger *et al.*, 2019). PacBio has developed a tool for this purpose (Biosciences, 2019b), which combines multiple subreads from the same ZMW using a statistical model to produce one highly accurate consensus sequence. However, this is a computationally intensive process and requires multiple passes for error correction to work reliably (Biosciences, 2019b). Considering the high accuracy of the latest PacBio Sequel chemistry (Rhoads and Au, 2015), the improvement in accuracy due to generating a consensus sequence is small compared to using the Read Subsampling tool. As the accuracy of raw PacBio sequence improves over time and the error rate between raw reads and consensus sequence decreases the runtime advantage of using the Read Subsampling tool relative to generating a consensus sequence, becomes more significant. When appropriate, subsampling longest subreads per CLR using *SequelTools* will reduce redundancy and data size.

Random CLR subsampling, another function of *SequelTools*’ Read Subsampling tool, will further reduce data size to any size desired. This could be useful for bootstrapping or generating a test data set for the purpose of developing or testing tools or computational pipelines. This could also be useful for situations where using large data sets is not feasible, like in phylogenetics. Random CLR subsampling using the read subsampling tool from *SequelTools* allows the user to generate a subsample of their raw sequence data in an unbiased way that retains all information associated with one CLR across subreads and, potentially, scraps files.

### 3.3 The Read Filtering Tool

*SequelTools* also provides a read filtering tool, which would be very useful for those wishing to normalize their data. Read filtering can be done by minimum CLR length, having at least one complete pass of the DNA molecule through the polymerase, and/or having normal scraps adapters. Some users may wish to filter by minimum CLR length. We believe that the large majority of users that are using scraps reads will find value from filtering by the number of complete passes and normal scraps adapters. Scraps reads alone are affected by filtering by the number of complete passes and normal scraps adapters, because subreads data are, by default, set to a number of passes of one and a ZMW classification annotation of “normal” in the PacBio SAM format (Biosciences, 2019a). Filtering by normal scraps adapters refers to a ZMW classification annotation of “normal”, as opposed to “control”, “malformed”, or “sentinel”, and a scrap region-type annotation of “adapter”, as opposed to “barcode” or “LQRegion” (Biosciences, 2019a). When filtering by minimum CLR length, both scraps and subreads files are affected. Keep in mind that the calculated CLR length will include the merged data from both subreads and scraps files given to *SequelTools*. We believe that the Read Filtering tool will be valuable to the majority of people working with PacBio Sequel data who are working with scraps files due to the need to normalize data before performing downstream analysis.

### 3.4 Benchmarking

Each tool within *SequelTools*’ was tested and benchmarked on Condo, a High-Performance Computing Cluster, at Iowa State University, running the Red Hat Linux operating system. Varying number of CPU’s (Central Processing Units) (4 to 16, with increments of 1) and 8 SMRTCells from the PacBio reads of the NC358 maize genome (Bioproject ID XXXXXXXXXX, Biosample ID XXXXXXXXXX) (Ou *et al.*, 2019) was used for benchmarking. For the QC tool, both with and without scraps modes were benchmarked with fixed memory. All default QC tool options were used. The UNIX time command was used to collect the ‘real’ usage time for each run. On average it took less than 30 minutes to run with only scraps and little over an hour with both scraps and subreads (Figure 3). The runtime of the quality control tool is therefore tightly correlated with total input file size. However, using a greater number of cores did not affect the runtime with or without scraps files. This is probably due to using the same amount of memory for each run or the processing time is disk read/write bound. We also noticed similar behavior for Samtools, which is the most time intensive component of the QC tool (Figure 3).

**Figure 3:**
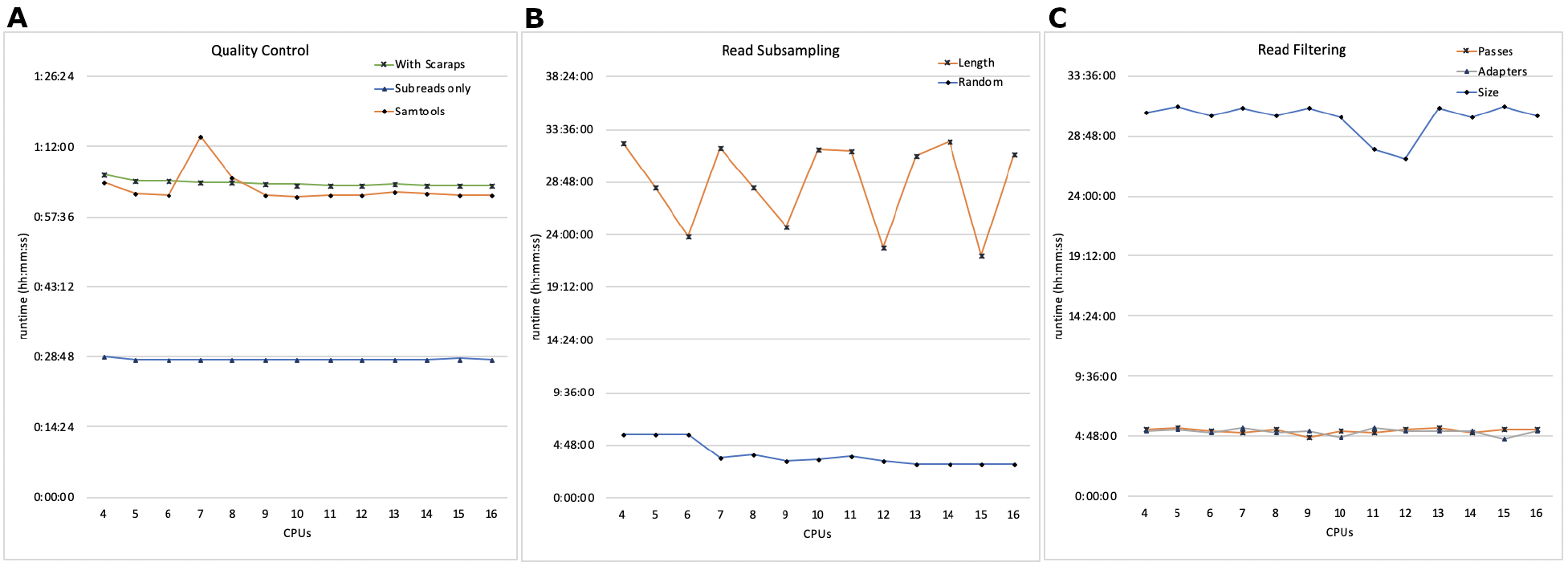
Runtime plot. The real runtime of eight SMRTcells of sequence from our benchmarking data set using between four and sixteen CPUs for all three of *SequelTools*’ tools: A) Quality Control, B) Read Subsampling, and C) Read Filtering. Benchmarking was done on Condo, an HPC cluster at Iowa State University.

**Figure 4:**
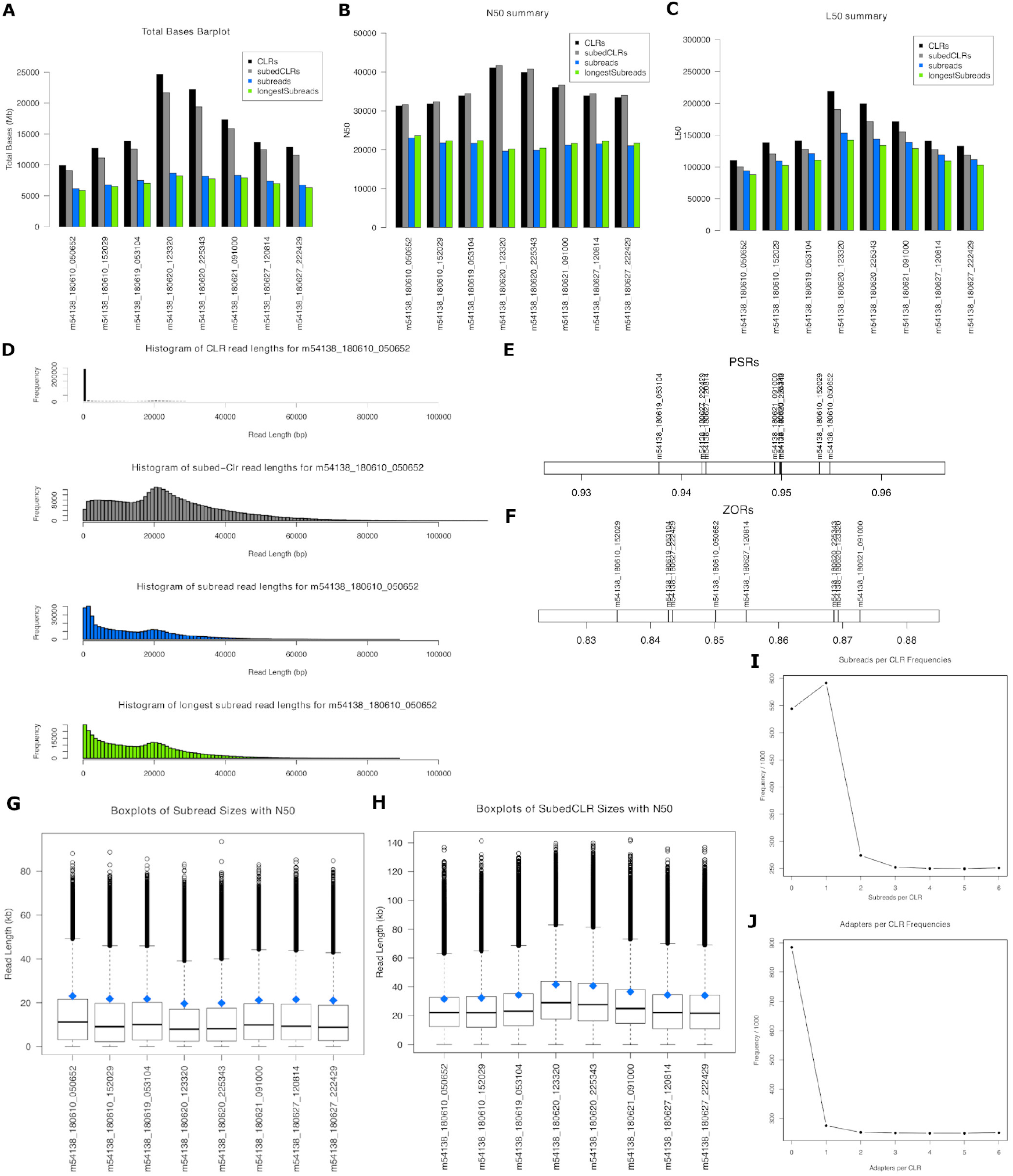
All plots generated by *SequelTools*. (A) A barplot of the sum of read lengths, (B) A barplot of N50s, (C) A barplot of L50s, (D) Read length histograms for m54138 180610 050652, (E) A two dimensional plot of PSRs, (F) A two dimensional plot of ZORs, (G) Boxplots of subread lengths with N50s as blue diamonds, (H) Boxplots of subedCLR lengths with N50s as blue diamonds, (I) A frequency plot showing the distribution of subreads per CLR, and (J) A frequency plot showing the distribution of adapters per CLR, for all SMRTcells in our benchmarking data set.

For the Read Subsampling tool, both random subsampling and subsampling for longest subreads was benchmarked with 4 to 16 processors with the same sized input data (8 SMRTcells). Subsampling by random CLRs took less than 6 hours with few processors but the runtime decreased steadily with the higher numbers of processors (∼3 hrs with 16 processors) (Figure 3). However, for subsampling for longest subreads, we did not find any improvements in runtime with additional CPUs. The bulk of the processing for subsampling for longest subreads is done serially via a Python script and additional CPUs will not help speed up this process. The 8 SMRTcell data, amounting to more than 130Gb and containing 4 million reads, takes about 32 hours for subsampling for longest subreads regardless of number of CPUs used (Figure 3).

For the Read Filtering tool, all three sub-tools were tested with similar benchmarking runs. Filtering by both normal scraps adapters and by number of passes takes about 5 hours to complete (Figure 3). Again, since the bulk of the time is used for actual read filtering serially via a Python script, the number of CPUs does not improve runtime for this tool. For length-based filtering, the minimum CLR length threshold of 1Kb was used and read filtering was performed on all 8 SMRTcells. This process took about 30 hours to complete (Figure 3). All three of *SequelTools*’ tools were run with both subreads and scraps files. The total size of this benchmarking data set is 277Gb.

## 4 Conclusion

*SequelTools* is an easy-to-install-and-use program that provides a variety of utilities for working with Pacbio Sequel raw sequence data including quality control, read subsampling, and read filtering. The QC tool calculates key statistics and generates publication-quality plots, providing all standard metrics for overall sequence quality including N50, read length and count statistics, PSR, and ZOR. *SequelTools*’ QC tool can evaluate eight SMRTcells from NC358 maize genome (Bioproject ID XXXXXXXXXX, Biosample ID XXXXXXXXXX) (Ou *et al.*, 2019) in about thirty minutes with subreads alone and in about an hour with the addition of scraps reads on our High-Performance Computing Cluster. Other than the proprietary PacBio *SMRTlink* program, which is time intensive, does not produce downloadable plots, and requires the user to set up a web server to install, there is currently no program available to compute these statistics. In addition to the QC tool, *SequelTools* has read subsampling and read filtering functions which allow the user to reduce their data size and to normalize their data, respectively. The author is not aware of any other program that provides these additional functionalities for PacBio Sequel data, except the *bamsieve* tool from SMRT Link (Biosciences, 2019c) which does random CLR subsampling but does not include scraps reads or longest subread subsampling. We therefore conclude that *SequelTools* is the only reasonable choice for quality control, the best choice for read subsampling, and the only choice for read filtering for users of PacBio Sequel sequencing data. We believe *SequelTools* will therefore contribute to the expansion of the use of PacBio’s highly valuable, and already popular, Sequel sequencing system.

## 5 Availability and Requirements

**Project name**: SequelTools

**Project home page**: https://github.com/ISUgenomics/SequelTools

**Operating systems**: Platform independent

**Programming languages**: Bash, Python, and R

**Other requirements**: samtools, awk

**License**: GNU GPL v3.0

## 6 Abbreviations

BAM: binary alignment/map format
CLR: continuous long read
CPU: Central Processing Unit
PDF: portable document format
PSR: polymerase-to-subread ratio
QC: Quality Control
RAM: Random-access memory
SAM: sequence alignment/map format
SMRT: Single Molecule Real Time sequencing technolog
ZMW: zero-mode waveguide
ZOR: zero-mode waveguide occupancy ratio

## 7 Acknowledgements

We would like to thank Maggie Woodhouse for proving comments on this manuscript, as well as Jim Coyle, director of HPC at Iowa State University and Andrew Severin at the Genome Informatics Facility for supporting the development of this tool.

## 8 Funding

This work was supported by the National Science Foundation Plant Genome Research Program Grant number IOS-1744001 and Principle Investigator Kelly Dawe.

## 9 Availability of data and materials

The test data set used for benchmarking is available in the NCBI repository, BioProject XXXXXXXXXX

## 10 Authors’ contributions

D.E.H. designed, implemented and maintained the code, and coordinated the research activity. D.E.H., M.B.H., and A.S. formed the project goals including suggesting new features, and contributed to data acquisition and visualization. M.B.H. and A.S. provided oversight and mentorship. A.S. performed bench-marking. M.B.H. acquired financial support for the project. D.E.H. and A.S. tested code components, wrote the manuscript, and designed the methodology. M.B.H. edited the manuscript. All authors read and approved the final manuscript.

## 11 Ethics approval and consent to participate

Not applicable.

## 12 Consent for publication

Not applicable.

## 13 Competing interests

The authors declare that they have no competing interests.

